# Mechanisms of surface and volume light scattering from *Caenorhabditis elegans* revealed by angle-resolved measurements

**DOI:** 10.1101/2025.09.24.678386

**Authors:** Zihao (John) Li, Christopher Fang-Yen

## Abstract

The use of the model roundworm *Caenorhabditis elegans* in has enabled pioneering discoveries in biology in part due to its optical accessibility. Optical imaging of *C. elegans* primarily relies on light scattering to generate contrast. However, the mechanisms of light scattering by *C. elegans*, and how they influence image contrast, are not well understood. We measured scattering and image contrast from *C. elegans* under varied illumination angles, wavelength, substrate composition, bacterial seeding, and index matching conditions. Combining experiments with ray-tracing simulations, we show that scattering arises from two interacting components: volume scattering within the body and surface scattering at interfaces between the worm and its surroundings. Surface scattering dominates at low angles and enhances animal boundaries, whereas volume scattering becomes more prominent at higher angles and improves visibility of internal structures. We identify practical conditions that optimize image contrast, including low-angle off-axis illumination, longer-wavelength (red) light, and low-scattering substrate media. Our work establishes a framework for understanding light scattering from small animals.

## INTRODUCTION

The microscopic nematode *C. elegans* is one of the most widely studied model organisms in biology due to its short life cycle, genetic accessibility, and compact anatomy.^1,2^ Its optical accessibility, fully mapped connectome,^3^ and rich repertoire of behaviors^4^ have made it a valuable system for *in vivo* imaging of neural activity,^5–7^ behavioral dynamics,^8–10^ and genetic perturbations.^10,11^ Recent advances in automation^11–16^ have enabled high-throughput genetic and pharmacological screening using *C. elegans*.

One of the strengths of *C. elegans* is its amenability to optical imaging. Many assays of behavior, morphology, aging, and development rely on obtaining images of worms with high contrast relative to their surroundings.^17–21^ Since like most biological tissues, *C. elegans* exhibits relatively low optical absorption over optical wavelengths, most of the optical contrast is provided by scattering, i.e. changes in direction of incident light rays.^22^

In any imaging system, lenses or other optical elements are configured to focus light collected from a certain range of angles. Two imaging modes can be differentiated by whether the illumination angles are within the range of collection by the optical system.^23,24^ In a bright field imaging configuration, the illumination rays are within the collection angle of the optics. Optical scattering by the object diverts some of the light away from the collection angles, causing the object to appear darker than its surroundings. By contrast, in a dark field imaging configuration, the illumination light is outside the collection angle of the optical system; scattering by the object causes some of this light to be collected, and the object appears bright against a dark background.

The amount and angular distribution of light scattering are highly sensitive to illumination angle, wavelength, and the optical properties of the object and background medium. In practice, many imaging systems rely on empirically chosen parameters, without a systematic understanding of how these variables affect image contrast.

Light scattering in biological tissues has previously been studied primarily in the context of biomedical imaging.^22,25^ Prior theoretical descriptions of light scattering in biological tissues include those applying Mie theory^26,27^ which describes scattering from spheres as a function of diameter and refractive index, and the Henyey–Greenstein phase function,^28,29^ which describes the degree of anisotropy occurring during scattering. Numerical simulations using the discrete dipole approximation and discrete source method have been used to model light scattered by red blood cells.^30,31^ Angle-dependent scattering profiles have been characterized for tissue sections,^32–34^ tissue-mimicking phantoms,^35^ and single cells.^34,36^ Cell structure has been shown to be an important factor modulating optical scattering from unicellular marine organisms.^37^ However, to our knowledge mechanisms of light scattering from intact multicellular living organisms with specific geometries, such as *C. elegans*, have not been examined in detail.

Although high-resolution imaging methods such as Differential Interference Contrast (DIC) and fluorescence microscopy are widely used for detailed anatomical phenotyping, many *C. elegans* assays, such as high-throughput screening^16,18,19,38^ and behavioral tracking,^4,10,39^ rely on label-free bright field or dark field imaging, where image contrast largely depends on optical scattering from the animals. Therefore, establishing a mechanistic and quantitative description of how intact *C. elegans* scatters light as a function of various imaging parameters is important for image optimization.

In this study, we sought to understand the mechanisms of light scattering from *C. elegans* and explore the imaging conditions that optimize optical contrast. We first quantify scattering functions and image contrast as a function of illumination angle, revealing a pronounced contrast maximum at around 12°. We then test the wavelength dependence of scattering and show that longer wavelengths increase image contrast. To explain these empirical trends, we proposed a model that optical scattering from *C. elegans* arises from a combination of volume scattering within the animal body and surface scattering due to the refractive index mismatch at the interface between the worm and its surroundings. Next, we perform empirical index-matching analysis and computational ray tracing to verify our model. Our computational model successfully reproduces the angular and spatial distribution of optical scattering observed from experimental measurements. We find that surface scattering dominates at low angles whereas volume scattering becomes more prominent at higher angles. Furthermore, we demonstrate how these mechanisms translate to practical choices of illumination in *C. elegans* microscopy: low-angle illumination produces images emphasizing body boundaries whereas high-angle illumination produces images that better resolve internal anatomy. Finally, we test how substrate composition affects image contrast. We find that compared with conventional substrates based on agar, optical contrast is much higher on substrates based on low-scattering biopolymers gelatin and gellan gum.

Our work provides a mechanistic framework for understanding light scattering in intact organisms and offers practical guidance for optimizing quality in *C. elegans* imaging. Moreover, the physical principles outlined here will inform optimization strategies for imaging other organisms.

## RESULTS

### Angle dependence of scattering functions and image contrast

To establish a quantitative description of light scattering and optical contrast from *C. elegans*, we first measured scattering magnitude and image contrast as a function of illumination angle.

We first illuminated worms (N2) on agar plates at specific angles θ, using a ring of light-emitting diodes (LEDs) and captured images using a camera and a lens (Fig. 1A(i), Materials and Methods). Using a ring of LEDs creates azimuthally uniform illumination. We set the collection angle of the system to be low (numerical aperture, NA = 0.031) to optimize measurements of scattering angle (Fig. 1A(ii), Discussion) at the cost of lower spatial resolution.

**Fig. 1.**
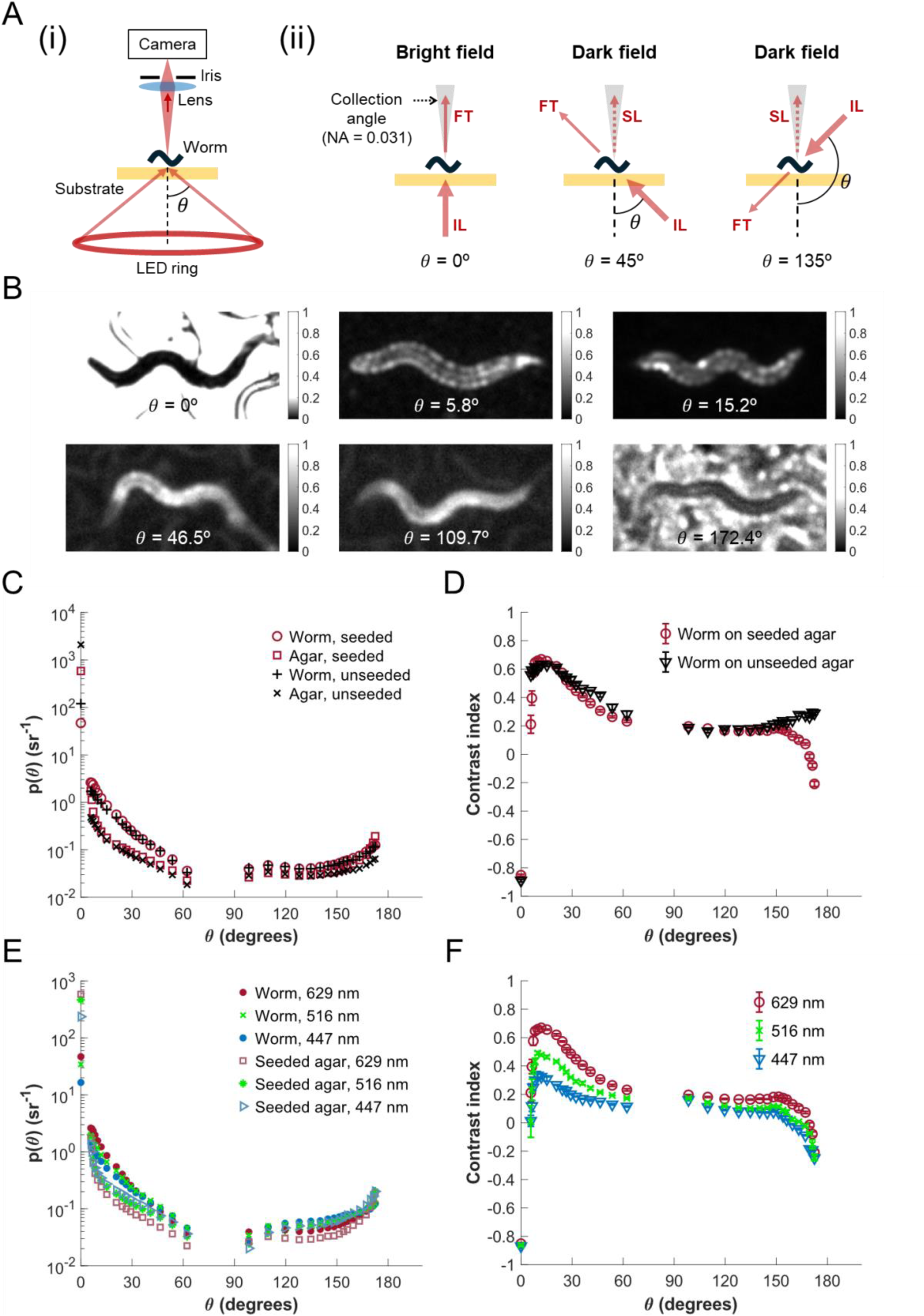
Angle-resolved, wavelength-dependent scattering functions and image contrast for *C. elegans* on agar substrate. (A) (i) Experimental setup, showing illumination applied at specific polar angles θ. (ii) Detection geometry and imaging modes for different illumination angles θ. FT: forward transmission; IL: illumination; SL: scattered light. The illumination is applied across all azimuthal angles. (B) Representative images of worms and background at different illumination angles (FOV: 1.11 × 0.56 mm). (C) Scattering functions for *C. elegans* on seeded and unseeded agar substrate under red light illumination. Maximum relative standard error (RSE) is 14% for seeded agar and 9% for all other categories. (D) Image contrast as a function of illumination angle θ for *C. elegans* on seeded and unseeded agar plate under red light illumination. (E) Scattering functions for *C. elegans* on seeded agar plates under red, green, and blue light illumination. Maximum RSE is 9%, 6%, 12% for worms, and 14%, 22%, 11% for seeded agar, under red, green, and blue illumination. (F) Image contrast index for *C. elegans* on seeded agar as a function of illumination angle and wavelength. For (D) and (F), the error bars represent standard error. For (C) and (E), error bars are not shown due to the high density of the datapoints and the small RSE. Statistical details are provided in Table S1. For (C)-(F), each datapoint represents the mean of measurements across N = 3 to 5 animals. For (B)-(F), all the animals are *C. elegans* N2 at day-1 adulthood.

Fig. 1A(ii) depicts the geometries of illumination and detection for representative θ values. It also demonstrates how the image mode changes with θ. For θ near 0°, the illumination rays are nearly vertical, and the light collected by the optics consists of mainly forward transmission. The worm scatters portion of the light out of the collection angle of the optics, causing the animal to appear darker than the background. As a result, for θ near 0°, the animal is imaged in a bright field mode, in which the background appears brighter than the worm. For larger values of θ, for example 45°, the illumination rays form an oblique angle with the vertical optical axis. The forward transmission is outside of the collection angle and only the portion of the light scattered by the worm can be collected by the optics. In this scenario, the animal appears brighter than the background and the imaging is in a dark field mode. For θ larger than 90°, for example 135°, the illumination is at the same side of the lens and the camera. In this case, only the light scattered backward by the worm can be collected by the optics and therefore the animal is also imaged in dark field.

As described above, under illumination at variable θ angles, animals on agar seeded with food bacteria can be imaged in either bright field or dark field mode (Fig. 1B). We also found that *C. elegans* exhibits distinct spatially inhomogeneous scattering patterns across different illumination angles (Fig. 1B). The contrast of the animal against its background also changes with θ (Fig. 1B).

Using the images recorded with different illumination angles, we calculated the scattering function p(θ), which has units of sr^-1^ (inverse steradian) and describes the probability per solid angle of a photon scattering by a polar angle θ relative to its original trajectory (Fig. 1A).^40,41^ Note that the scattering function p(θ) measured using the setup depicted in Fig. 1A(i) is azimuthally symmetric because the illumination is applied across all azimuthal angles. The value of the scattering function at θ = 0° represents the probability of a ray to be transmitted without scattering.

We measured scattering functions under red light illumination (peak wavelength 629 nm) for *C. elegans,* agar, and bacteria-coated (seeded) agar (Fig. 1C). For both *C. elegans* and agar backgrounds, the scattering function is highest at θ = 0°, decreases from 0° to 62° (*p* < 0.005), and increases modestly from 128° to 172° (*p* < 0.005).

To determine the effect of scattering angle θ on image contrast, we defined a contrast index equal to the difference in irradiance between worm and background, both measured at the camera, divided by the sum of irradiance of the worm and background (see Materials and Methods). We calculated the contrast index as a function of θ (Fig. 1D). For θ = 0°, the contrast indices are negative for worms on both seeded and unseeded agar plates (*p* < 0.001). This is because at θ = 0°, the system is in the bright field regime (Fig. 1A(ii)), in which the worm appears darker than its surroundings. For θ > 0°, in the dark field imaging regime (Fig. 1A(ii)), the contrast index increases sharply with θ and reach a maximum at 12° (*p* < 0.001, Fig. 1D). For θ close to 180°, we observe a negative contrast index for worms on seeded plates (*p* < 0.001, Fig. 1D). These results suggest that optimal image contrast can be obtained by applying illumination around 12° relative to the optical axis.

### Wavelength dependence of scattering functions and image contrast

Having found that illumination angle is an important modulator of optical scattering and contrast, we next asked whether other illumination settings also play important roles. We reasoned that scattering from *C. elegans* would vary with illumination wavelength, since optical scattering from biological tissue typically shows a strong wavelength dependence.^42,43^

To determine the wavelength dependence, we measured the angle-resolved scattering functions for *C. elegans* on seeded agar plates under red (peak wavelength 629 nm), green (516 nm), and blue (447 nm) LED illumination (Fig. 1E). We found that the scattering functions for green and blue light show similar overall trends as that of red light.

For θ = 0°, the scattering functions of *C. elegans* are higher for red than green (*p* = 0.005), and higher for green than blue (*p* < 0.001) (Fig. 1E). It indicates that the proportion of the light that is forward transmitted without scattering increases with wavelength, consistent with results from prior studies that the optical reduced scattering coefficient 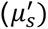 generally decreases with wavelength.^42^

We also found that illumination wavelength has a prominent effect on the image contrast (Fig. 1F). The peak contrast indices for red illumination are higher than green ( *p* < 0.001), and green higher than blue (*p* = 0.009). These results suggest that red illumination is better than green or blue in terms of generating contrast in images.

### Empirical analysis of volume and surface scattering

The empirical measurements of scattering from *C. elegans* led us to ask what mechanisms contribute to the observed angle dependence of the scattering functions. We reasoned that light scattering from *C. elegans* could arise from two sources: (1) volume scattering within the animal body due to refractive index variations and (2) surface scattering from refractive index mismatch at the boundaries between the worm and the air and/or the substrate (Fig. 2A).

**Fig. 2.**
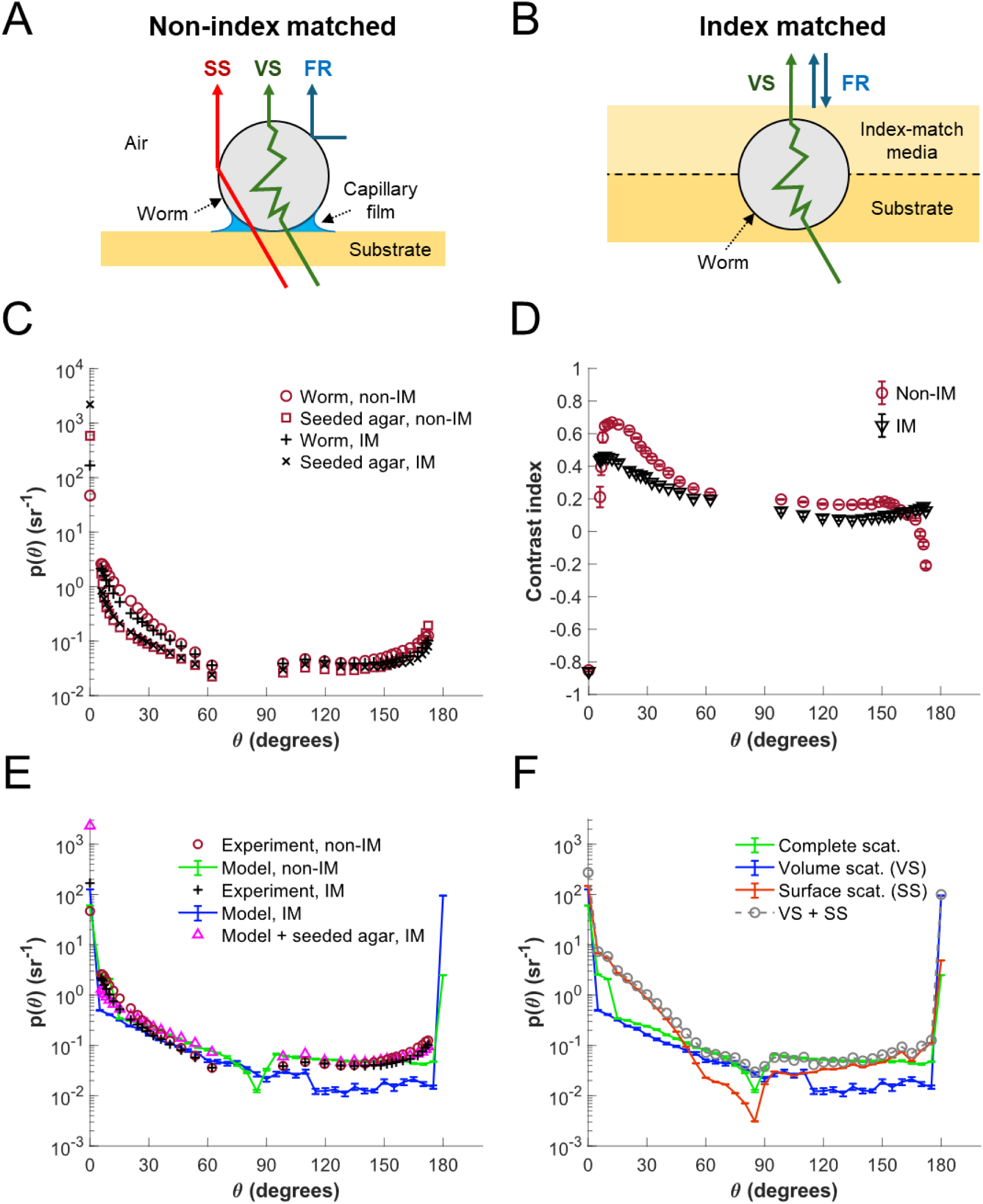
Analysis of volume and surface scattering from *C. elegans*. (A) Schematic of light scattering from *C. elegans* on a moist substrate. SS: surface scattering; VS: volume scattering; FR: Fresnel reflection. (B) Schematic of light scattering from *C. elegans* on the substrate underneath an index-matching media. (C) Angle-resolved scattering functions for *C. elegans* and seeded agar for the non-index matched (non-IM) and the index matched (IM) cases. Maximum RSE is 14% for all the categories. Error bars are not shown due to the high density of the datapoints and the small RSE. Statistical details are provided in Table S1. (D) Angle-resolved image contrast of *C. elegans* on seeded agar for non-IM and IM cases. Error bars represent standard error of the mean. (E) Simulated angle-resolved scattering functions of *C. elegans* obtained from the computational model for non-IM and IM cases, overlaid with the experimental results. (F) The simulated scattering functions for the volume scattering only, the surface scattering only, the sum of the volume and surface scattering, and the complete scattering. For (C) and (D), each datapoint represents the mean of measurements across N = 3-5 animals. For (E) and (F), datapoints and error bars represent the mean and standard error across N = 10 simulation trials. For (C) - (E), all the animals are *C. elegans* N2 at day-1 adulthood.

To help distinguish between these two types of scattering, we first isolated the volume scattering by minimizing the surface scattering via approximate refractive index matching. Most biological tissues have an index of refraction slightly higher than that of water.^44^ The *C. elegans* cuticle has been measured to have a refractive index about n = 1.35-1.37, only slightly higher than that of water.^45^ To achieve approximate index matching, we placed a gelatin pad (n ≈ 1.36) on the agar plate in order to eliminate the worm-to-air interface and surround the animals with media with similar refractive index to that of the worm (Fig. 2B). After index matching, the observed scattering function should be almost entirely due to volume scattering.

Fig. 2C depicts the scattering functions for *C. elegans* and bacteria-seeded agar in the index-matching experiment. For θ = 0°, scattering functions for the index-matched case are higher than for the non-index-matched case for both worm (*p* = 0.007) and background (*p* < 0.001), implying that index matching increases the forward transmission.

We calculated the contrast indices due to the volume scattering only, for the worm and background (Fig. 2D). For θ = 0°, in the bright field imaging regime, the image contrast for index matching is not significantly different to non-index matching (*p* = 0.753). For θ > 0°, in the dark field imaging regime, the image contrast reaches maximum around 10° for index matching. The peak contrast index for index matching is lower than non-index matching (*p* < 0.001, Fig. 2D), implying that surface scattering helps to increase image contrast.

As in the non-index matched case, we observed that volume scattering in *C. elegans* is dependent on wavelength (Fig. S1A). Besides, red light generates peak image contrast higher than green (*p* = 0.005), and green higher than blue (*p* = 0.032) (Fig. S1B). Taken together, our results show that optimal image contrast for *C. elegans* in an index-matched environment can be obtained by applying red illumination with an angle of 10°-15° relative to the optical axis.

These findings inform useful guidelines for optimizing image quality for *C. elegans* in liquid-based assays^19,46,47^ or other high-index environments.

### Mechanisms of volume and surface scattering

The experimental observations of optical scattering in the non-index matched and index matched scenarios lead us to pose several questions. What mechanisms contribute to the angular dependence of the optical scattering from *C. elegans* and how does it shape the nonlinear change of the image contrast (Fig. 2C and D)? What is the interplay between the volume and the surface scattering and how do they contribute to the overall scattering? Why does the optical scattering exhibit spatially inhomogeneous patterns across the worm’s body, sometimes appearing to highlight the outline of the animal (Fig. 1B)?

To explore the mechanisms underlying the angle-dependent scattering, we created a simple computational model of the worm and its surroundings (Fig. 2A). We modeled the adult *C. elegans* as a cylinder (diameter = 50 µm, length = 1000 µm, n = 1.33) on the surface of a rectangular cuboid (n = 1.33), representing the substrate. A concave shaped film (n = 1.33) formed at the either side of the cylinder (Fig. 2A and Fig. S2), representing the capillary water film that forms near the points of contact between the worm and the agar substrate.^48^ We set the volume scattering within the cylinder to follow the Henyey-Greenstein phase function,^28^ which is widely used to model light scattering within biological tissues.^29^ The anisotropy parameter g is set to 0.83 and the scattering mean free path set to 12 µm (optimization of free parameters is described in Materials and Methods). To simulate the index matched case, we eliminated the worm-to-air interface by adding a medium (n = 1.33), representing the index matching pad, to the model (Fig. 2B).

Using these models, we simulated scattering functions for both the index matched and non-index matched cases. We observed that the simulated scattering functions show broad agreement with the experimental observations for both the index matched and non-index matched cases (Fig. 2E). For θ = 0°, simulated scattering function for index matching is higher than non-index matching (*p* < 0.001), suggesting that index matching increases the forward transmission, which is consistent with the experimental observation (Fig. 2C and E). The simulated scattering functions are highest at θ = 0°, decrease from 0° to 65° (*p* < 0.001), and increase from 150° to 180° (*p* < 0.001), showing an overall trend broadly consistent with the experimental results (Fig. 2E). We also observed that the index-matching model does not fit the experimental data well for θ between 90° and 180°. We attribute this discrepancy to scattering from the bacteria-seeded agar, which is not included in the simulation. To account for this effect, we summed the index-matching model and the experimental measurements of seeded agar (Fig. 2C). This improved the agreement between the model and the experimental data for θ between 90° and 180° (Fig. 2E).

In the simulation, the back scattering detected at θ = 180° (Fig. 2E) is due to Fresnel reflection (i.e. that occurring at interfaces between media of different refractive index) (Fig. 2A and B).^49^ In the index matched case, the flat horizontal upper surface of the index-matching medium (Fig. 2B) creates a Fresnel reflection collected by the detector at θ = 180° (Fig. 2E). The Fresnel reflection created by the convex surface of the simulated worm (Fig. 2A) in the non-index matched case is weaker than that of the index matched scenario (*p* < 0.001, Fig. 2E).

In addition to calculating the scattering due to volume scattering only (via index matching), we calculated the scattering due to surface scattering alone (by removing volume scattering), the sum of the volume only and surface scattering only models, and finally the scattering in the complete model incorporating both volume and surface scattering (Fig. 2F).

In the surface scattering only model, the worm and water film act as an approximately cylindrical lens. Due to the lensing effect, the surface scattering dominates over the volume scattering from 0° to 50° (*p* < 0.01), but weaker than the volume scattering for from 55° to 85 ° (*p* < 0.001), as shown in Fig. 2F. Due to the Fresnel reflection, the surface scattering is stronger than the volume scattering from 115° to 175° (*p* < 0.001, Fig. 2F).

By comparing the sum of the volume only and surface scattering only models with the complete model incorporating both volume and surface scattering, we can assess the degree of interaction between volume and surface scattering. If volume and surface scattering occur independently, we would expect the sum of the volume only and surface scattering only to approximately equal the complete model. This is indeed the case for large angles, from about 60° to 150° (Fig. 2F), because Fresnel reflection from the upper worm surface is independent of volume scattering occurring inside the animal. By contrast, for angles from 5° to 45°, the complete scattering is much smaller than the surface scattering only model (*p* < 0.001). This is because rays coming from below the worm must first go through volume scattering in the worm’s body and then surface scattering before being collected by the detector. In this case, the incident ray directions are largely randomized by the volume scattering, causing most of them to scatter into larger angles. On the other hand, for angles from 5° to 45°, the lensing effect of the surface scattering in the complete model helps refracting the rays to reach the detector, causing the complete scattering to be somewhat higher than that of the pure volume scattering (*p* < 0.02, Fig. 2F).

We asked how well other models could also describe our scattering data. Mie theory describes scattering from spheres as a function of diameter and refractive index and is a widely used model for studying light scattering from biological tissues.^26,27^ We applied Mie theory to simulate the scattering function for a sphere with volume, surface area, or cross-sectional area (Fig. S3) equivalent to that of an adult worm (diameter = 50 µm, length = 1000 µm, n = 1.33). We then compared the results from the three Mie models with the experimental data for worms on seeded agar (Fig. S3). We found that all three Mie models overestimate the direct transmission at 0° (*p* < 0.001) while underestimating scattering at angles greater than 90° (*p* < 0.005), except from 135° to 140° (Fig. S3). Compared to the three Mie models, our model shows better agreement with the empirical data (Fig. S3).

### Inhomogeneous spatial distribution of light scattering

Having found that volume and surface scattering collectively shape the angle dependence of scattering, we next asked whether these two components also contribute to the spatially inhomogeneous patterns observed in the worm images (Fig. 1B). First, we measured the cross-sectional scattering profile for the animal at various illumination angles (Fig. 3A-D). We noticed that in some images the worm appeared brightest along its edges, whereas in others it appeared brightest at its center. To quantify how much the light scattering is spatially located around the edges of the worm body compared to the center, we defined the edge-to-middle contrast index (EMCI) (see Materials and methods). The EMCI has a range from -1 to 1 with positive EMCI indicating the scattering is brighter around the edges than in the middle. We measured EMCI as a function of θ for the index matched and non-index matched cases (Fig. 3E and F).

**Fig. 3.**
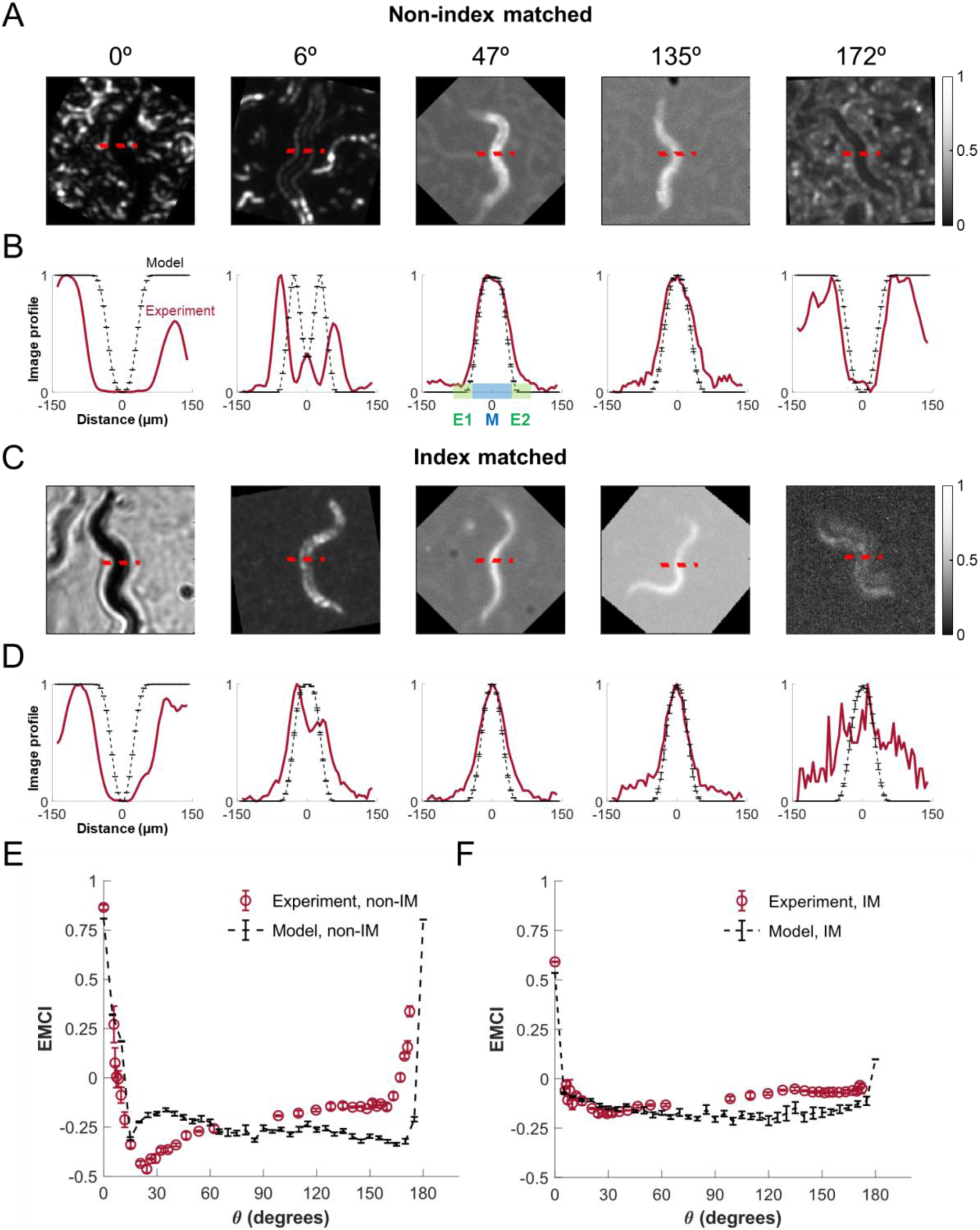
Spatial distribution of light scattering from *C. elegans*. (A) Images recorded at representative angles for *C. elegans* on seeded agar, without an index matching media (FOV = 1.11 × 1.11 mm). The red dashed line highlights the pixels used for calculating the cross-sectional scattering profiles at the midpoint of the animal. (B) Scattering profiles across the midpoint of the animal for individual representative angles in the non-index matched case shown in (A). E1, M, and E2 represent the first edge portion, the middle portion, and the second edge portion of the profile used for calculating EMCI. (C) Images recorded at representative angles for *C. elegans* on seeded agar underneath an index matching media. (D) Scattering profiles across the midpoint of the animal for individual representative angles in the index matched case shown in (C). (E) Experimental and simulated EMCIs as functions of θ for the non-index matched (non-IM) scenario. (F) Experimental and simulated EMCIs as functions of θ for the index matched (IM) scenario. For (B), (D), (E), and (F), datapoints and error bars for the simulation data represent the mean and standard error of the mean, respectively, across N = 10 simulation trials. For (E) and (F), datapoints and error bars for the experimental results represent the mean and standard error across N = 3 animals. All animals are *C. elegans* N2 at day-1 adulthood.

In the non-index matched case, for θ < 6°, stronger scattering is experimentally observed around the edges and weaker in the middle, yielding positive EMCI (*p* < 0.05, Fig. 3E). For θ from 10° to 165°, scattering is weaker around the edges and stronger in the middle, yielding negative EMCI (*p* < 0.02). When θ > 168°, the EMCI turns positive again (*p* < 0.02). By contrast, the EMCI with index-matching is not significantly higher than zero (*p* > 0.80) for all the experimentally tested θ except for 0° (Fig. 3F). These results indicate that the high scattering observed around the edges for θ close to 0° and 180° for the non-index matched scenario are due to the surface scattering. In comparison, the volume scattering consistently concentrates around the middle of the animal body with different θ.

Next, we ask whether the observed spatial distribution of scattering can be reproduced by our computational model depicted in Fig. 2A and B. The profiles of scattering across the simulated worm for the representative θ in the scenarios of non-index matching and index matching are shown in Fig. 3B and D, respectively. The simulated EMCIs as functions of θ for both scenarios are respectively shown in Fig. 3E and F. The results from our simulation show broad agreement with the experimental data, further supporting the validity of our computational model.

Combining the results from Fig. 2F, we can describe the mechanisms underlying the angle-dependent spatial distribution of the scattering. For θ = 0°, in the bright field imaging regime, the light incident to the animal is scattered out of the collection angle of the detector (Fig. 1A(ii)), causing the animal to appear darker than the background (Fig. 1B, Fig. 3A-D), yielding negative image contrast (Fig. 1D and Fig. 2D) and positive EMCI (Fig. 3E and F). For small θ > 0°, in the dark field imaging regime (Fig. 1A(ii)), the surface scattering is dominant (Fig. 2F) and contributes to the brightened boundaries of the animal due to the lensing effect, yielding positive EMCI (Fig. 3B and E). For larger scattering angles, the volume scattering is dominant (Fig. 2F), and the scattering spatially distributes around the center, yielding negative EMCI (Fig. 3E and F). The negative contrast index and the positive EMCI observed near θ = 180° for worm on bacteria-seeded agar (Fig. 1B, D, F, Fig. 3A, B, and E) is due to the Fresnel reflection from the substrate.

These findings hold important implications for choosing optimal illumination angles for different purposes. For some applications where animal tracking and the identification of small independent objects (such as embryos and larvae) are more important, smaller angles would be preferable since the surface scattering highlights the object boundaries and overall contrast is highest. For cases in which resolving the internal structures of *C. elegans* is more important, the illumination should be set to higher angles where the volume scattering is dominant.

### Optimal illumination for microscopy

Having identified that volume scattering is dominant at high angles whereas surface scattering is more prominent at low angles and spatially concentrated around edges, we next asked how these findings can translate to practical guidelines to form optimal illumination for microscopy of *C. elegans*. For this purpose, we acquired images of worms using imaging systems with higher magnification and NA (Materials and methods), under various angles of illumination (Fig. 4).

**Fig. 4.**
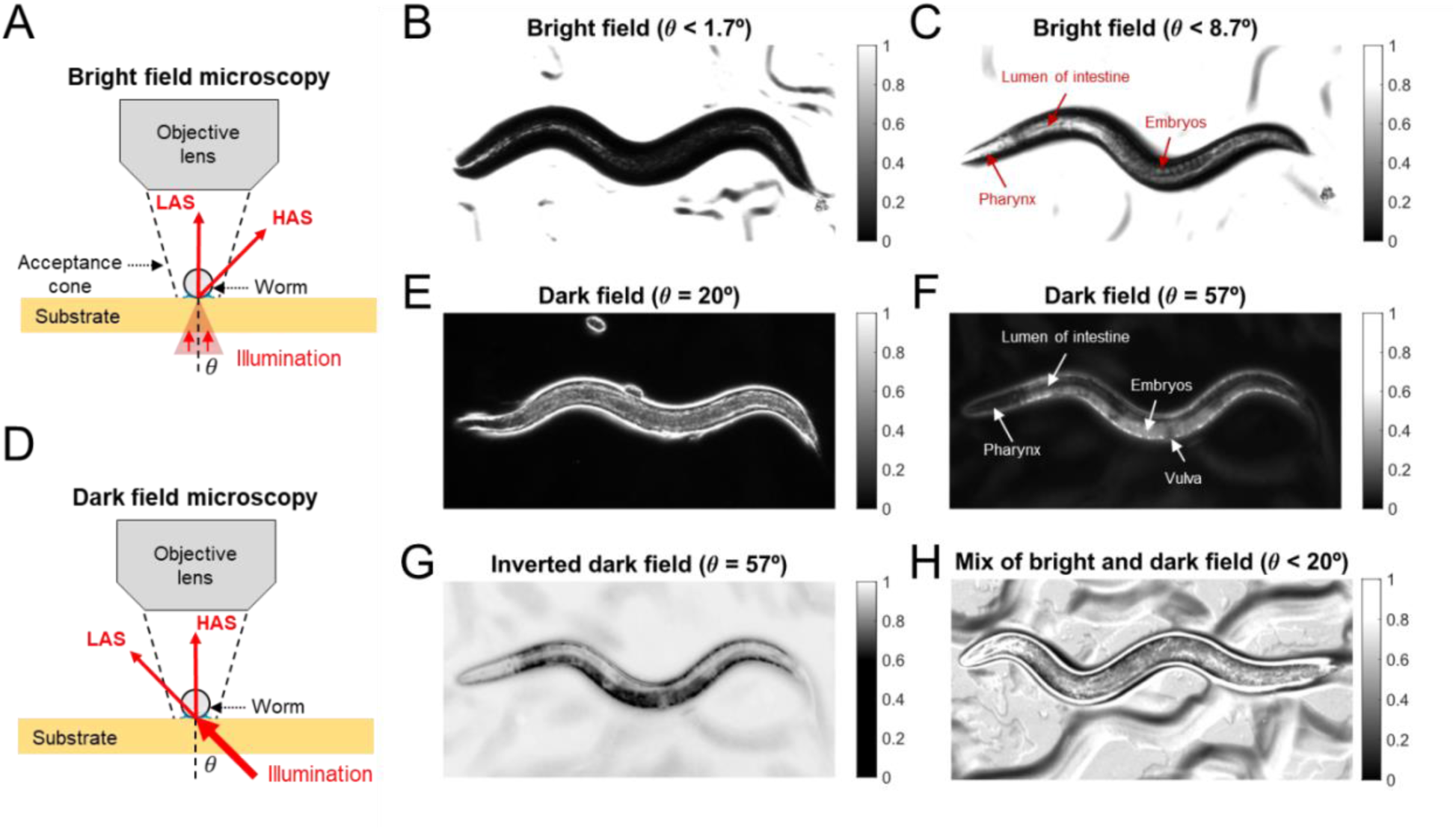
Effects of illumination angle on bright field and dark field microscopy. (A) Schematic diagram of bright field microscopy. LAS: low-angle scattering; HAS: high-angle scattering. (B) A bright field image of *C. elegans* under narrow-angle illumination (θ < 1.7°) FOV: 1.0 x 0.5 mm. (C) A bright field image of *C. elegans* under wider-angle illumination (θ < 8.7°). (D) Schematic diagram of dark field microscopy. (E) A dark field image of *C. elegans* under low-angle illumination (θ = 20°). (F) A dark field image of *C. elegans* under higher-angle illumination (θ = 57°). (G) Computationally inverted version of F yields an image similar to bright field illumination. (H) A combination of bright field and dark field images of *C. elegans* under illumination applied both inside and outside of the collection angle range of the objective lens (θ < 20°). The NA of the objective lens is 0.3 for (B) and (C), and 0.14 for (E) – (H). All the animals shown in this figure are *C. elegans* N2 at day-1 adulthood.

In bright field microscopy (Fig. 4A), the angular range of illumination θ is smaller than that collected by the objective lens. Some of the light incident on the worm scatters out of the collection angles, causing the worm to appear darker than its background. When θ is small, the light collected from the animal is dominated by low-angle scattering (Fig. 4A), which is largely contributed by surface scattering (Fig. 2F). The surface scattering helps improve the contrast of the edges with a tradeoff of lower resolution for the internal structures (Fig. 4B), consistent with the results from our spatial analysis (Fig. 3A, B, and E). To better visualize internal features, the angular range of illumination should be larger so that more light scattered to high angles due to volume scattering can be collected (Fig. 4C). Compared to an image using narrow angle illumination (Fig. 4B), wider-angle illumination better resolves internal structures within the body, such as pharynx, intestine, and embryos (Fig. 4C).

In dark field microscopy (Fig. 4D), the illumination is outside the acceptance angle of the objective lens. Optical scattering diverts some of the light into the acceptance angle, and the worm appears bright against a dark background (Fig. 4E-F). When the illumination angle is low, more low-angle scattering (surface scattering) is collected, contributing to bright edges (Figs. 3A, B, E, and 4E) and high contrast against the background (Figs. 2D and 4E). In contrast, larger θ causes more high-angle scattering (volume scattering) to be collected, which facilitates the visualization of internal structures (Fig. 4F). Compared to Fig. 4E, higher-angle illumination helps resolving the internal organs (Fig. 4F and G).

For both bright field and dark field imaging, it is important for the illumination angles to be either inside or outside of the lens’ acceptance range, but not both simultaneously. Illuminating with a range of angles both inside or outside of the lens’ acceptance range results in a combination of bright field and dark field images, resulting in low image contrast (Fig. 4H).

Important heuristics can be translated from our findings to practical guidelines for common microscopy in *C. elegans* labs. First, for observations of animals through stereoscopes or high-magnification microscopes, it is preferential to use transillumination (θ < 90°) as compared to epi-illumination (θ > 90°), because image contrast under epi-illumination is lower (Figs. 1D, F, and 2D).

Second, illumination angles need to be tailored for different purposes. For animal tracking and identification of embryos and larvae, illumination can be set to lower NA for highlighting object boundaries to obtain maximum contrast. To achieve this on a stereoscope, the angle of the reflective mirror under the sample stage can be adjusted so that the illumination light rays are nearly parallel to the optical axis. If available, the size of the illumination aperture can be reduced. For commercial high-magnification microscopes, where Köhler illumination^50^ is commonly used, the illumination NA can be lowered by reducing the size of the condenser aperture. In contrast, for visualization of internal structures of *C. elegans*, illumination needs to be set to higher NA.

### Substrate-dependent image contrast

In addition to illumination geometry and wavelength, image contrast is influenced by the amount of scattering in the substrate background. While *C. elegans* is commonly cultured on agar media, other gelling agents may also be used instead of agar.^9,51,52^ We asked how the composition of the substrate might affect image contrast. We prepared three substrates including agar, gelatin, and gellan gum (Materials and methods), at concentrations that yielded similar gel strengths, and measured scattering functions and image contrast for animals on these substrates (Fig. 5).

**Fig. 5.**
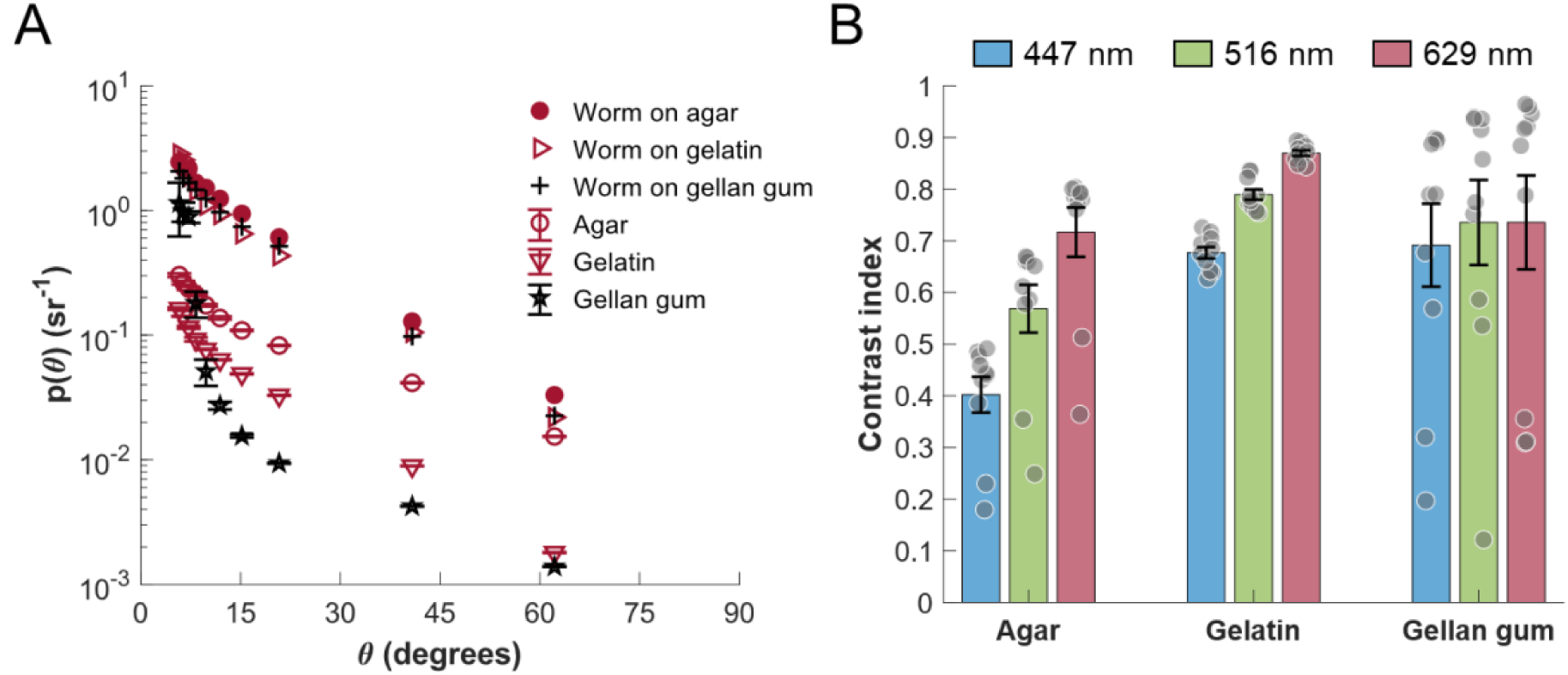
Scattering functions and image contrast for different substrate polymers. (A) Angle-resolved scattering functions for *C. elegans* on unseeded media made with agar, gelatin, and gellan gum, under red illumination. Each datapoint and error bar represents the mean and standard error across N = 3 to 7 animals. Maximum RSE for worms are 6.8% on agar, 7.3% on gelatin, and 8.0% on gellan gum. Some error bars are not shown due to the high density of the datapoints and the small RSE. Statistical details are provided in Table S1. (B) Image contrast for *C. elegans* on unseeded agar, gelatin, and gellan gum under red, green, and blue illumination. Each dot represents the mean of measurements across N = 3 to 7 animals for an individual illumination angle. Error bars represent the standard error across N = 10 illumination angles. All the animals are *C. elegans* N2 at day-1 adulthood.

Our results show that the scattering functions for all three substrates decrease with θ from 6° to 62° (*p* < 0.01) but show different magnitudes (Fig. 5A). Overall scattering from gelatin (*p* < 0.005) and gellan gum (*p* < 0.005) is smaller than from agar for θ from 10° to 62°. Lower background scattering results in the peak image contrast for animals on gelatin (*p* < 0.001) and gellan gum (*p* < 0.001) higher than agar (Fig. 5B).

We observed that the scattering functions for all the substrates are dependent on illumination wavelength (Fig. S4) and longer-wavelength illumination generates higher peak image contrast (*p* < 0.05, Figs. 5B and S4).

We also notice that the imprint due to animal’s movement causes significant scattering on gellan gum for θ < 10° (Fig. S4F), which leads to reduced image contrast (Figs. 5B and S4F). These imprints may not be favorable for animal identification or tracking but could be helpful for studying locomotion trajectories.

## DISCUSSION

In this work, we investigated the mechanisms of light scattering from the nematode *C. elegans* using angle-resolved measurements and computational modeling. Our findings reveal that the light scattering from *C. elegans* arises from two primary sources: volume scattering within the animal body and surface scattering at the interfaces between the animal, air, and the moist substrate. These two types of scattering exhibit distinct angular and spatial profiles and contribute differently to the overall scattering. Surface scattering is more prominent at small and large angles, while volume scattering dominates at intermediate angles (Fig. 2). Volume scattering is highest at the central region of the animal (Fig. 3) and helps resolve internal anatomical structures (Fig. 4). On the other hand, surface scattering enhances the animal’s boundaries (Fig. 3), which may aid animal detection and tracking but obscure internal features (Fig. 4). The interplay between the volume and surface scattering suggests that the optimal imaging configuration should be tailored to the specific applications, for example, behavioral tracking versus anatomical visualization.

We found that the image contrast is highly sensitive to multiple parameters, including the illumination angle, wavelength, and the type of imaging substrates. These findings inform a set of imaging parameters that enhance image contrast. The simple strategy we provided here does not require specialized optics and labels, and therefore it is readily adaptable to many systems, for example, microscopy,^17,53–56^ high-throughput screening setups,^10,18,19,38,39^ microfluidic chambers,^46,57,58^ and automation systems.^12,38,59,60^

Together, this study provides a biophysical understanding for the light–tissue interactions in multiple-scattering organisms and offers physically informed strategies for optimizing the image quality in a wide range of imaging systems. Our findings hold immediate value for researchers using *C. elegans* as a model system, and broader relevance for biological imaging across disciplines, such as genetics, neuroscience, and pharmacology.

### Limitations of the study

It is important to acknowledge the limitations of our work. First, some sources of error may affect the accuracy of the scattering function measurements. For example, the imperfect alignment of the components to the optical axis would also contribute to the discrepancy in the measurements of the angle. The LEDs, the FOV of the camera, and the iris of the lens have non-zero sizes, such that the scattered lights we collected are from a finite range of angles. Therefore, in order to minimize the angular uncertainty, we selected lights for particular angles by intentionally setting the NA of the lens to be low (F-number f/16, NA = 0.031), with an acceptable cost of lower spatial resolution of the images (Fig. 1B, Fig. 3A and C).

There are a few factors that may affect the agreement of our computational model to the experimental data. First, the animal is modeled as a simple cylinder, where the geometry of head and tail and the bending of the body are not captured. Second, optical absorption from the animal is not modeled. Third, we did not model the scattering due to the substrate or bacteria, causing the background noise underestimated in the detected signal. Next, the optical simulation is based on geometric ray tracing where the wave properties of the light, such as diffraction, interference, and polarization, are not modeled.

This study focuses on a single developmental stage (day-1 adult) of a specific *C. elegans* strain (N2). Variations in body morphology, internal structures, or cuticle properties across life stages or genetic backgrounds may alter light scattering patterns.^17^ Future work could explore how scattering profiles change across developmental stages and in genetic mutants with altered anatomy.

The calculation of the scattering function and image contrast is on an individual-animal basis, which may neglect the variability between different organs and tissues. For example, pharyngeal muscles are less scattering compared to embryos or intestine (Fig. 4F). Future work will investigate the scattering profiles for different anatomical structures.

We expect the mechanisms of light scattering we identified here to have better explanatory power for transparent, absorption-minimal biological samples, such as other nematodes,^61^ fruit fly larvae,^62^ zebrafish larvae,^63^ and human organoids.^64^

## RESOURCE AVAILABILITY

## Data and code availability

The data and code used in this study are available from https://github.com/Zihao-celegans/Worm-Scattering, including:

- Dataset S1: Raw data for this paper.
- Design File S1: Design files (in STEP format) for the 3D models representing *C. elegans* and its surroundings, generated using Autodesk Fusion.
- Software S1: Source files (in ZMX and ZPL format) for conducting the optical simulations using non-sequential mode in Ansys Zemax OpticStudio 2024 R2.
- Software S2: Source MATLAB scripts for all data analysis in this study.

## ACKNOWLEDGMENTS

This research was supported by The National Institutes of Health (1R01DA056358). The *C. elegans* N2 strain was provided by the *Caenorhabditis* Genetics Center, funded by the NIH Office of Research Infrastructure Programs (P40 OD010440). We thank Anthony Fouad, Animesh Biswas, and Hongfei Ji for helpful discussions and suggestions.

## AUTHOR CONTRIBUTIONS

Conceptualization, Z.J.L. and C.F.-Y.; methodology, Z.J.L. and C.F.-Y.; investigation, Z.J.L.; writing—original draft, Z.J.L.; writing—review and editing, Z.J.L. and C.F.-Y.; funding acquisition, C.F.-Y.; resources, C.F.-Y.; supervision, C.F.-Y.

## DECLARATION OF INTERESTS

The authors declare no competing interests.

## METHODS

## KEY RESOURCES TABLE

| REAGENT OR RESOURCE | SOURCE | IDENTIFIER |
| --- | --- | --- |
| Experimental models: Organisms/strains |  |  |
| *Caenorhabditis elegans*/N2 | *Caenorhabditis* Genetics Center | https://cgc.umn.edu/strain/N2 |
| *Escherichia coli*/OP50 | *Caenorhabditis* Genetics Center | https://cgc.umn.edu/strain/OP50 |
| Software and algorithms |  |  |
| Fusion v.2605.1.52 | Autodesk | https://www.autodesk.com/products/fusion-360/overview |
| Zemax OpticStudio 2024 R2 | Ansys | https://www.ansys.com/products/optics/ansys-zemax-opticstudio |
| MATLAB R2025b | MathWorks | https://www.mathworks.com/products/matlab.html |
| ImageJ 1.54g | ImageJ | https://imagej.net/ij/ |

## EXPERIMENTAL MODEL AND STUDY PARTICIPANT DETAILS

The experimental model used in this study was the N2 strain of *Caenorhabditis elegans*. Animals were maintained on nematode growth medium (NGM) plates seeded with the OP50 strain of *Escherichia coli* at 20°C using standard methods.^65^ The *C. elegans* and *E. coli* strains were provided by the *Caenorhabditis* Genetics Center. All the animals used in the experiments were hermaphrodites on the first day of adulthood. No additional institutional permission was required for work with *C. elegans* under local institutional policies.

## METHOD DETAILS

### Imaging system

Our imaging system contains a LED ring comprised of 36 RGB LEDs (Qasim, 95 mm diameter), a 3D-printed culture plate holder, a camera lens (Nikon Micro-NIKKOR, effective focal length 105 mm, set to f/16, with C-mount adapter) and a CMOS camera (Imaging Source, DMK 33GP031) (Fig. 1A). All optical elements are mounted to a 1.5” diameter, vertically oriented post (Thorlabs P14) through post mounting clamps (Thorlabs C1511) and they are aligned to be centered about a common optical axis. The distance between the lens and the plate is 400 mm.

The illumination angle θ can be varied by translating the LED ring along the optical axis. The LED ring can be mounted below the agar plate to provide angles <90° or above the plate to provide angles between 90° and 180°. To obtain measurements at 0°, we positioned a single LED (3.5 mm × 3.5 mm size) a distance 465 mm directly under the plate. Due to the constraints of the optical mounts, some angles near 0°, 90°, and 180° were unavailable.

The LED array is programmable to emit red, green, or blue light. We measured the peak wavelengths of the LEDs using a fiber optic coupled spectrometer (Ocean Optics FLAME-S-XR1) to be 629, 516, and 447 nm. The spectral widths were approximately 16, 26, and 14 nm (FWHM), respectively.

### Measurement of scattering functions

The angle-resolved scattering function *p*(θ) describes the probability of a photon scattering into a unit solid angle oriented at an angle θ relative to its original trajectory.^40,41^ Here we calculated *p*(θ) via the expression (Eq. 1)

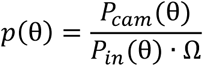

where *P*_*cam*_(θ) is the optical power received by the camera over the region of interest (ROI), *P*_*in*_ (θ) the total optical power incident on the object, and Ω the solid angle collected by the camera lens. We measured incident irradiance using an optical power meter (Coherent FieldMate Laser Power Meter with OP-2 VIS sensor). To calculate *P*_*cam*_(θ), we used the expression (Eq. 2)

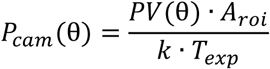

where *A*_*roi*_ is the area of the ROI, *T*_*exp*_ the exposure time, and *k* is the conversion coefficient from irradiance to pixel value. The measured *k* values for our camera in response to red, green, and blue light are 3.0, 3.3, and 3.5 × 10^5^ *ADU* · *mm*^2^ · *s*^−1^ · *μW*^−1^ respectively, where *ADU* is an analog-to-digital unit.

### Calculation of the contrast index

We defined the contrast index (*CI*) for an animal against its background as (Eq. 3)

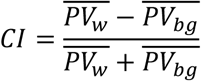

Where 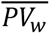 and 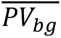 are the average pixel values inside the contour of the worm and the background. The contours of the animal were labeled using ImageJ (version 1.54g). The ROI for its background was chosen as a 1.1 mm x 1.1 mm rectangular region adjacent to the animal.

Since the scattering function *p*(θ) is proportional to the pixel values *PV*(θ) (Eq. 1 and 2), *CI* can be also expressed as (Eq. 4)

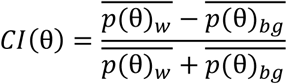

### Computational modeling

We conducted simulations using Ansys Zemax OpticStudio 2024 R2 optical design and analysis software in non-sequential mode. First, we constructed 3-dimensional models representing *C. elegans* on the culture substrate in the scenarios of non-index matching and index matching (Fig. 2A and B) using Autodesk Fusion (v.2605.1.52). The simulated animal is illuminated by 36 collimated light sources equally spaced in the azimuthal direction and with polar angle θ relative to the optical axis of the camera lens, which is modeled as a paraxial lens (f = 105 mm) with an iris (6.56 mm diameter) positioned 400 mm from the animal. The lens forms an image at an image distance of 142.37 mm. The image is captured by an incoherent irradiance detector. For each angle tested we performed 10 trials of ray tracing, with each trial containing 2 × 10^8^ analyzed rays. We recorded an irradiance map for each trial. Next, we calculated the scattering function *p*(θ) based on Eq. 1.

To model the volume scattering, we set the scattering within the simulated animal to follow the Henyey-Greenstein phase function^28^ (Fig. 2A and B). The two parameters, the anisotropy factor (g) and mean free path (MFP), were chosen by minimizing the mean squared error between the simulated scattering function and the experimental observations in the index-matched scenario (Fig. 2B and E). The optimized value for the anisotropy parameter was g = 0.83 and the mean free path was MFP = 12 µm.

After obtaining the optimized values for g and MFP, we simulated the complete scattering from the animal using the non-index matched model (Fig. 2A). We optimized the surface scattering by adjusting the shape of the capillary film to minimize the mean squared error between the simulated scattering function and the experimental data (Fig. 2E and Fig. S2).

### Edge-to-middle contrast index (EMCI)

To calculate the EMCI for the scattering profile across the animal (Fig. 3A-D), we first symmetrically divided the distribution into three portions (Fig. 3B), including the first edge portion (42 µm length), the middle portion (84 µm length), and the second edge portion (42 µm length). Next, we calculated the average value of the scattering function for both edge portions 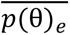 and for the middle portion 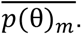 We then calculated EMCI as (Eq. 5)

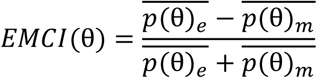

### Bright field and dark field microscopy

The bright field microscopy images of *C. elegans* (Fig. 4B and C) were obtained using an objective lens (NA = 0.3, Leica 1× Plan Apochromat) on a stereo microscope (Leica M165 FC). The angular range of illuminations was controlled by adjusting the aperture size of the illumination condenser.

The dark field microscopy images of *C. elegans* (Fig. 4E-H) were obtained using an objective lens (NA = 0.14, f = 40 mm, Mitutoyo 5X Plan Apochromat) together with a tube lens (f = 100 mm, Thorlabs TTL100-A) in a custom-built infinity-corrected microscope. Similar to Fig. 1A, the illumination angle is controlled by positioning an LED ring concentric with the optical axis at various distances to the animal.

### Preparation of imaging substrates

To prepare different culture substrates used for comparison of image contrast (Fig. 5), we dissolve powders of agar, gelatin, and gellan gum in NGM liquid at 100° C and wait until they solidify at room temperature. The concentration of the agar was set to 17 g/L according to the standard protocol.^65^ In order to obtain gelatin and gellan gum substrates suitable for maintaining *C. elegans*, we adjusted their concentrations such that the resultant substrates have comparable mechanical strengths as the standard 1.7% agar plate. We measured their mechanical strengths by a puncture test, where the minimum forces for an indenter (active area 19 mm^2^) causing fractures on the substrate were recorded by a precision balance (Mettler Toledo MS1602S). After these tests we set the concentration of gelatin to be 10.1% (average strength 79.9 ± 6.7 kPa, mean ± SD, N = 10 plates) and the gellan gum 1.125% (average strength 79.9 ± 5.7 kPa, N = 10), such that their mechanical strengths are comparable to the standard 1.7% agar plates (average strength 85.1 ± 9.3 kPa, N = 10).

### Preparation of *C. elegans*

The *C. elegans* N2 strain used in this study was obtained from the *Caenorhabditis* Genetics Center and cultivated at 20°C on NGM plates with *E. coli* OP50 using standard methods.^65^ We performed timed egg laying assays within a 4-hour window for synchronizing the ages of the animals. We used day-1 adults for all the experiments. All experiments were performed at ambient temperature (approximately 20 °C).

## QUANTIFICATION AND STATISTICAL ANALYSIS

We performed statistical analyses using MATLAB. All two-sample comparisons were conducted using Welch’s t-tests, under the assumption of unequal variances. Major claims are supported by *p*-values in the main text. The statistical details can be found in Results, figure legends, and Table S1.

## Supplemental information

**Fig. S1.**
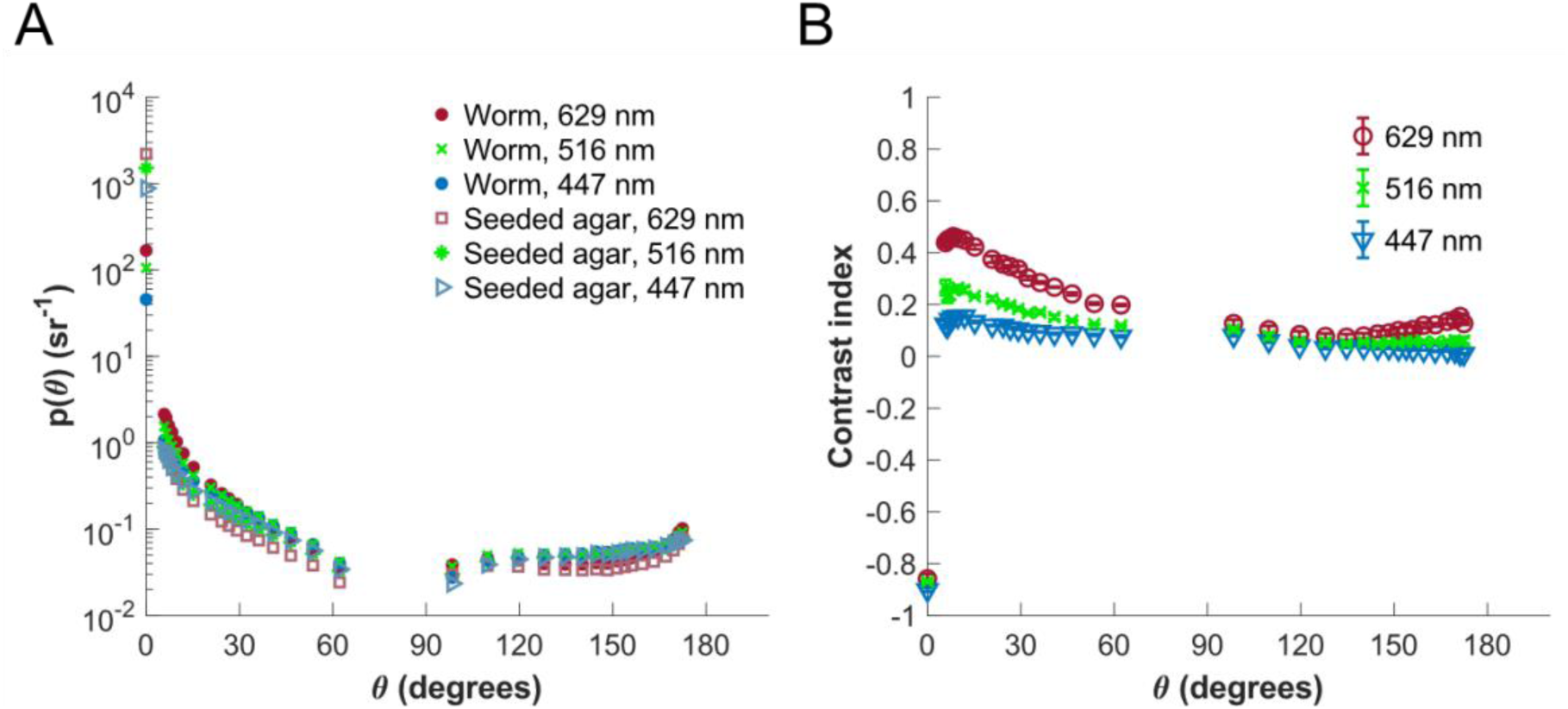
(A) Angle-resolved scattering functions for *C. elegans* and seeded agar for the index matched scenario under red, green, and blue illumination. Maximum relative standard error (RSE) is 14% for worms under red illumination and 7% for all other categories. Error bars are not shown due to the high density of the datapoints and the small RSE. Details of statistics are provided in Table S1. (B) Image contrast index for *C. elegans* on seeded agar as a function of illumination angle and wavelength for the index matched case. Error bars represent standard error. For both (A) and (B), each data point represents the mean of measurements across N = 3 to 4 animals. All the animals are *C. elegans* N2 at day-1 adulthood.

**Fig. S2.**
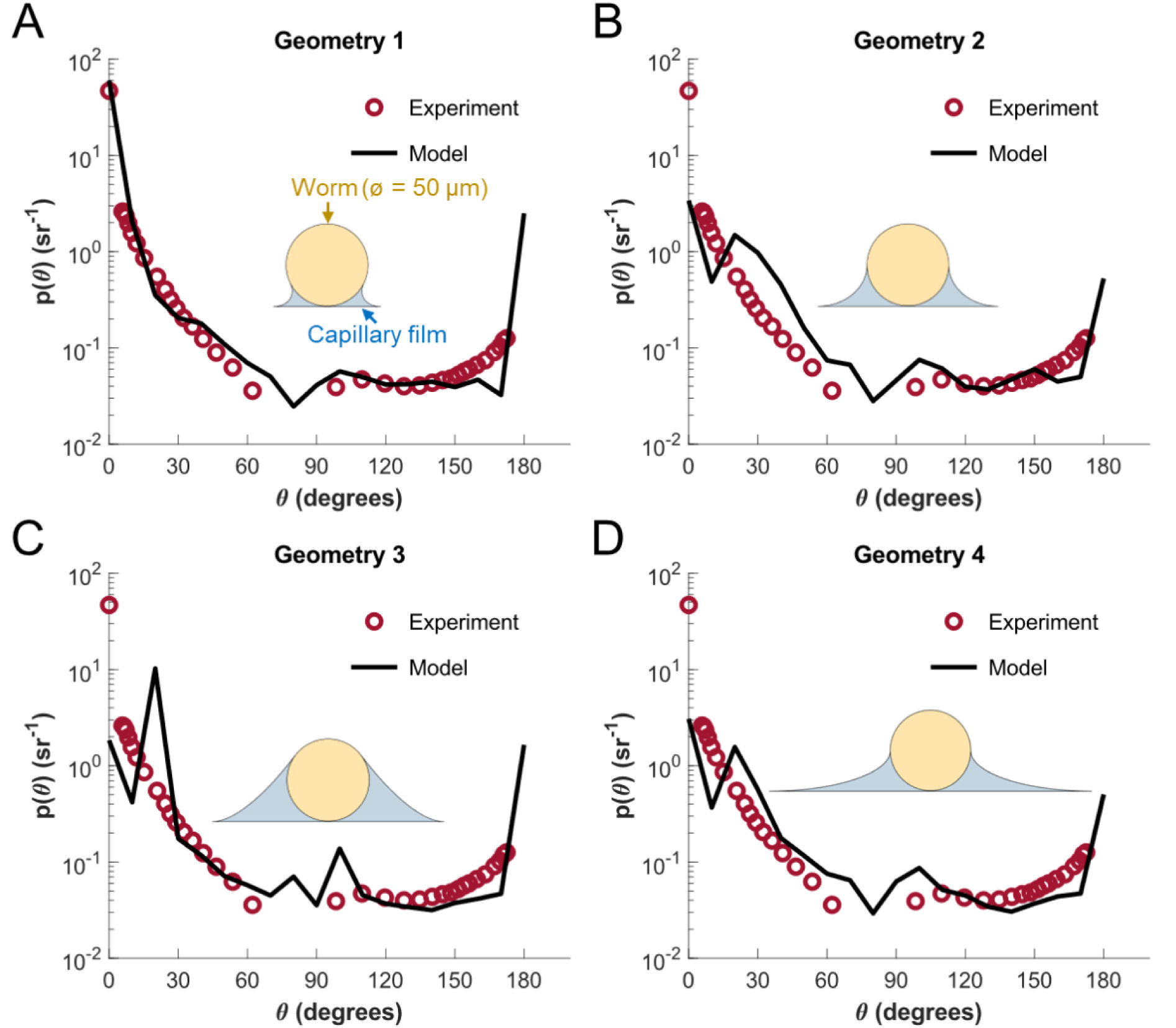
Scattering functions of *C. elegans* (N = 3 to 5 animals) obtained from the experiment (on seeded agar under 629-nm illumination) and the computational models with different geometries of capillary water film (N = 1 trial for each geometry). Insets: the cross section of the simulated animal and capillary water film. Each datapoint represents mean of measurements. Maximum RSE for the experimental data is 9%. All the animals are *C. elegans* N2 at day-1 adulthood.

**Fig. S3.**
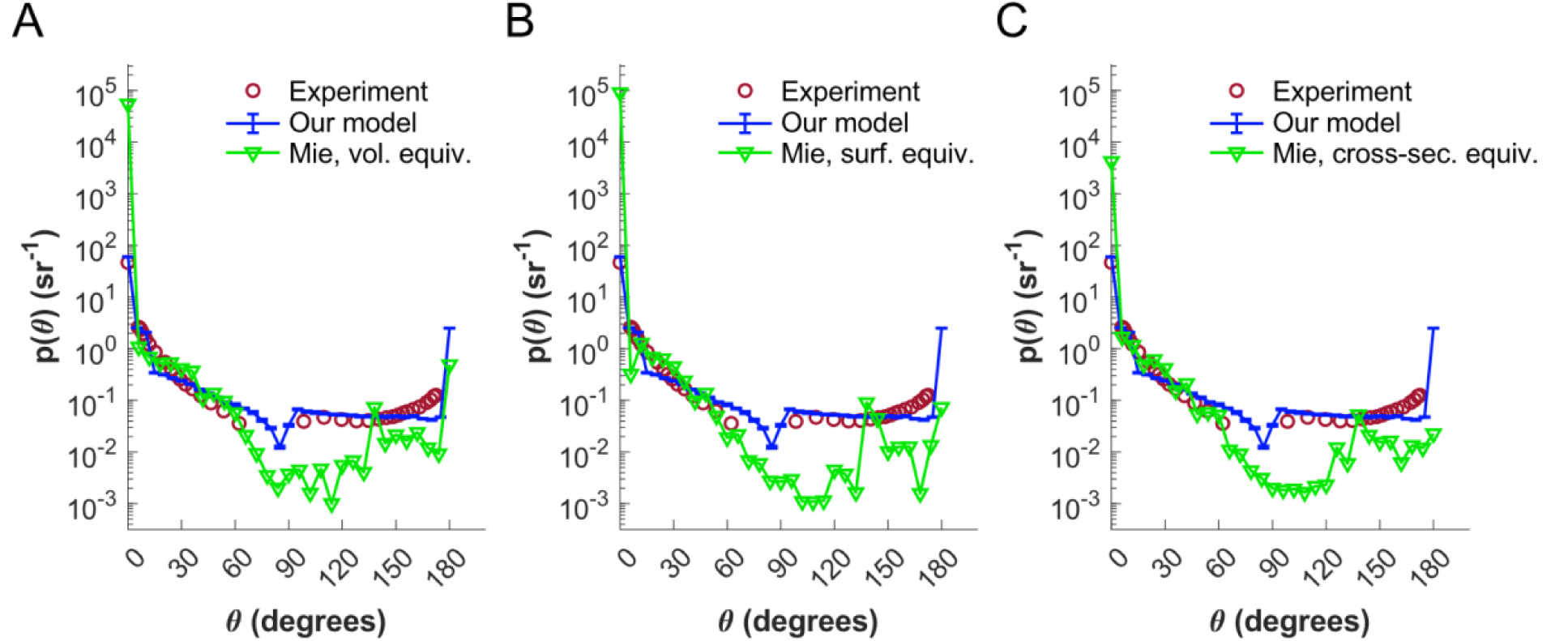
Scattering functions of day-1 adult N2 *C. elegans* (N = 3 to 5 animals) obtained from the experiment (on seeded agar under 629-nm illumination), our non-index matched model (N = 10 trials), and the Mie theory.^1^ Referencing to the dimensions of adult *C. elegans* (diameter = 50 µm, length = 1000 µm, n = 1.33), the Mie theory describes the scattering from a sphere with (A) equivalent volume (diameter = 155 µm), (B) surface area (diameter = 226 µm), or (C) cross-section area (diameter = 50 µm). Each datapoint represents mean of measurements. Maximum RSE for the experimental data is 9%. The error bars for the data from our model represent standard error.

**Fig. S4.**
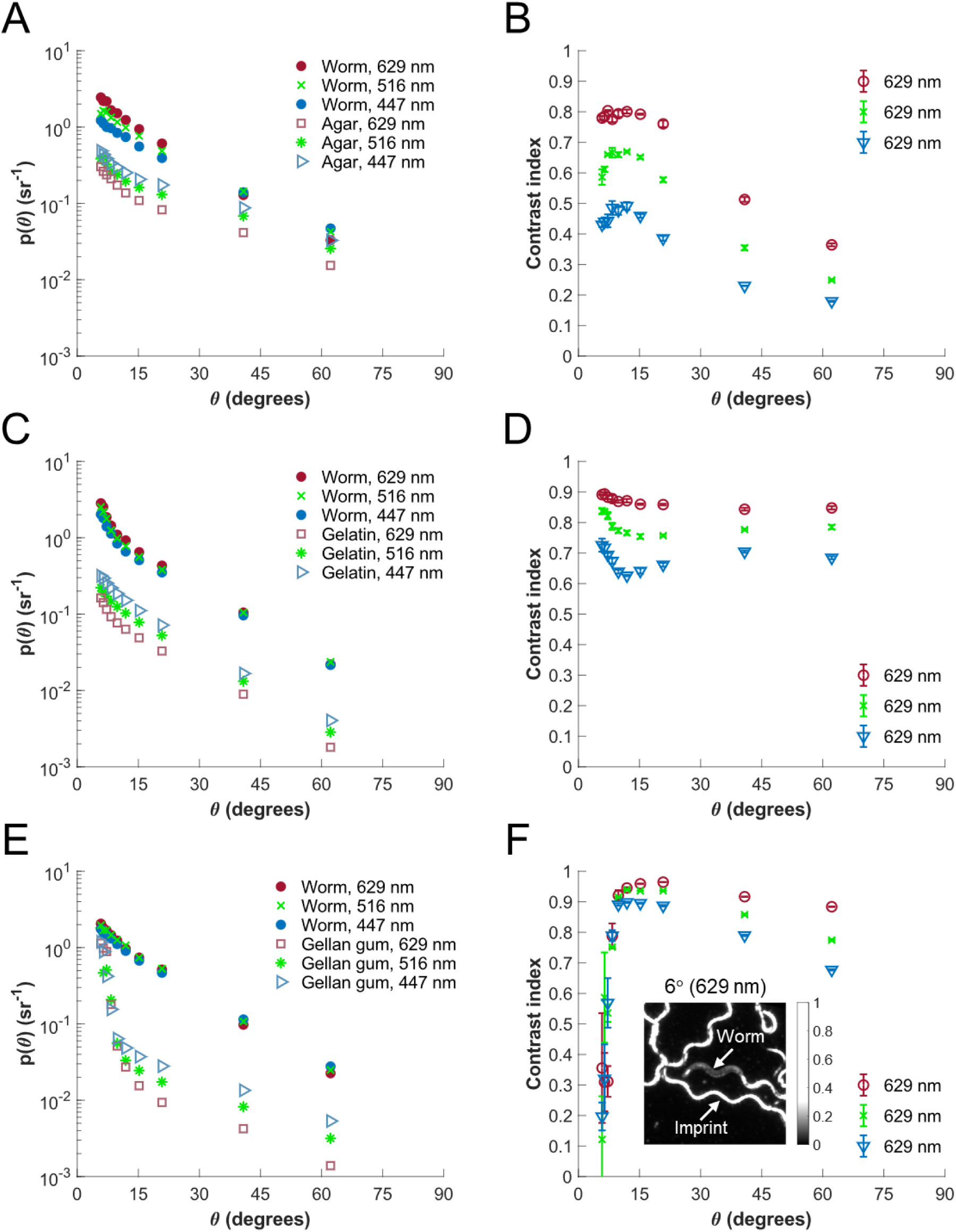
Angle-resolved, wavelength-dependent scattering functions and image contrast for *C. elegans* on unseeded media made with (A and B) agar, (C and D) gelatin, and (E and F) gellan gum. Inset of (F): a representative image of *C. elegans* and its imprints on unseeded gellan gum under a red illumination applied at 6° (FOV: 2.22 × 2.22 mm). For both (A) and (C), maximum RSE is 9% for all the categories. For (E), maximum RSE is 8% for worm and 46% for gellan gum. For (A), (C), and (E), error bars are not shown due to the high density of the datapoints, and statistical details are provided in Table S1. For (B), (D), and (F), error bars represent standard error. For all the panels, each data point represents mean of measurements across N = 3 to 7 animals. All the animals are *C. elegans* N2 at day-1 adulthood.

## Notes

### Competing Interest Statement

The authors have declared no competing interest.

### Summary of Updates

Various minor changes to the text and figures

