## Supplementary material for "Mechanisms of surface and volume light scattering from *Caenorhabditis elegans* revealed by angle-resolved measurements": Document S1

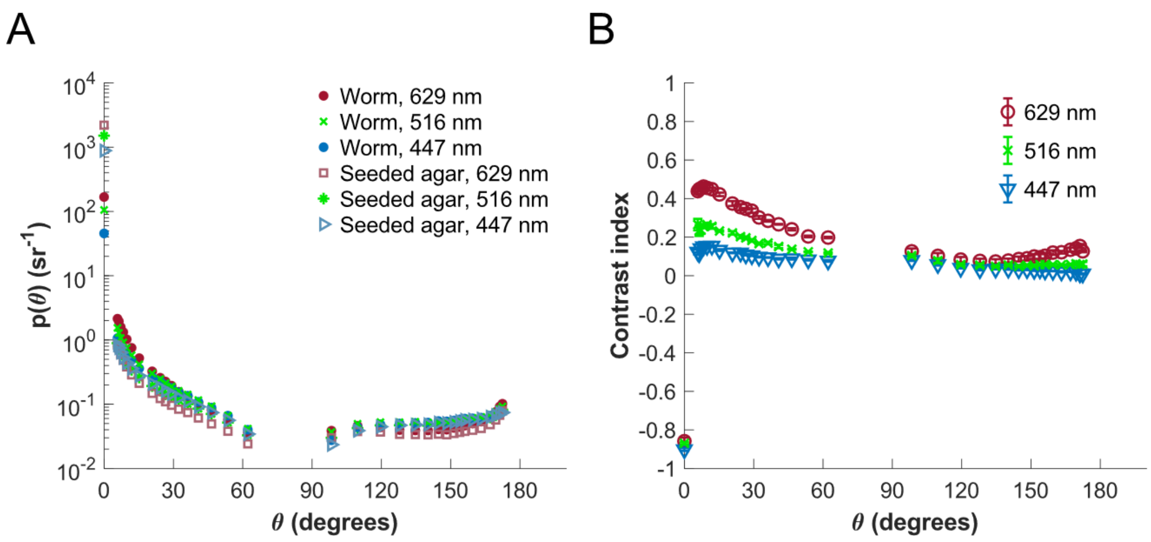


**Fig. S1.** (A) Angle-resolved scattering functions for *C. elegans* and seeded agar for the index matched scenario under red, green, and blue illumination. Maximum relative standard error (RSE) is 14% for worms under red illumination and 7% for all other categories. Error bars are not shown due to the high density of the datapoints and the small RSE. Details of statistics are provided in Table S1. (B) Image contrast index for *C. elegans* on seeded agar as a function of illumination angle and wavelength for the index matched case. Error bars represent standard error. For both (A) and (B), each data point represents the mean of measurements across N = 3 to 4 animals. All the animals are *C. elegans* N2 at day-1 adulthood.

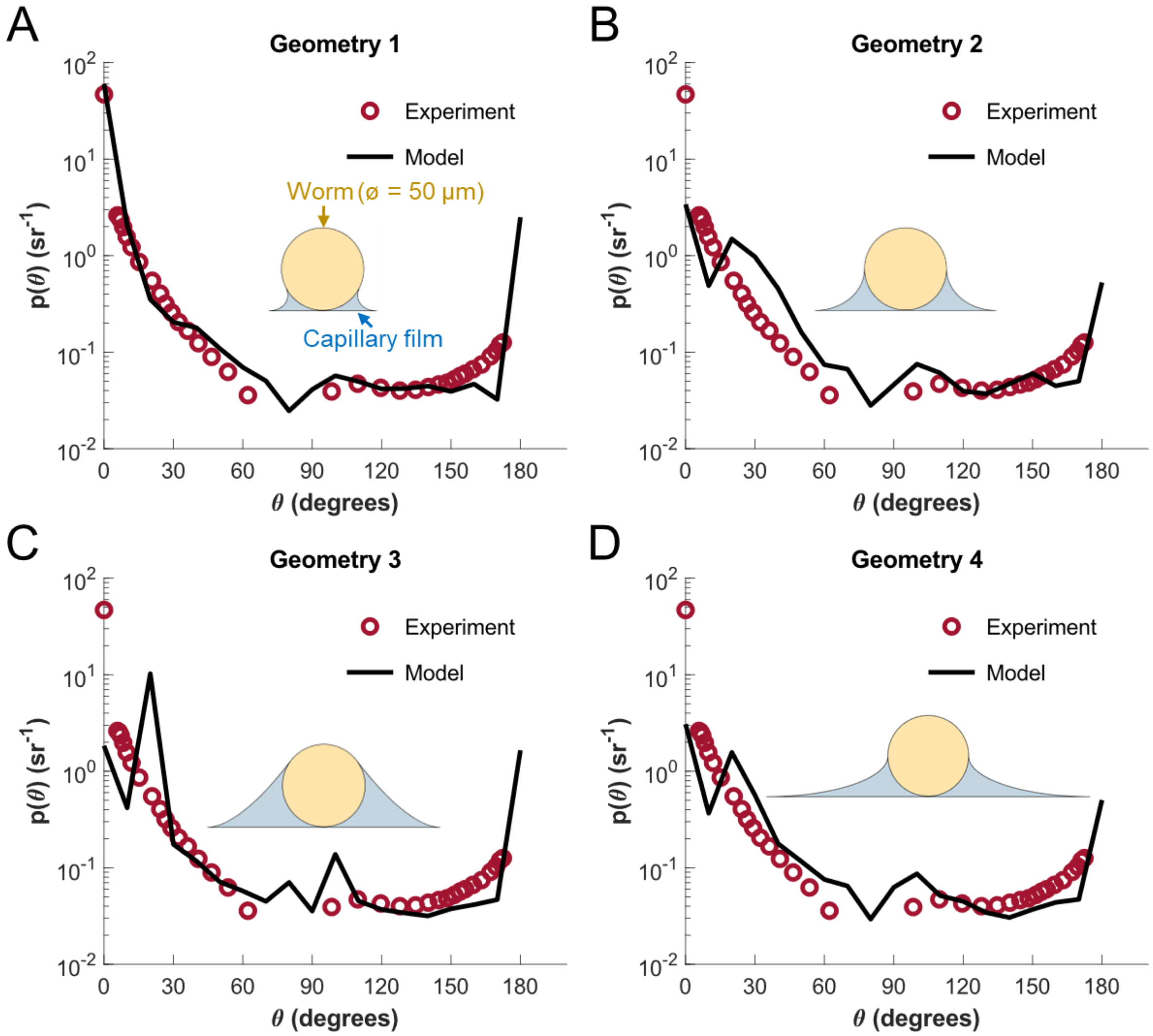


**Fig. S2.** Scattering functions of *C. elegans* (N = 3 to 5 animals) obtained from the experiment (on seeded agar under 629-nm illumination) and the computational models with different geometries of capillary water film (N = 1 trial for each geometry). Insets: the cross section of the simulated animal and capillary water film. Each datapoint represents mean of measurements. Maximum RSE for the experimental data is 9%. All the animals are *C. elegans* N2 at day-1 adulthood.

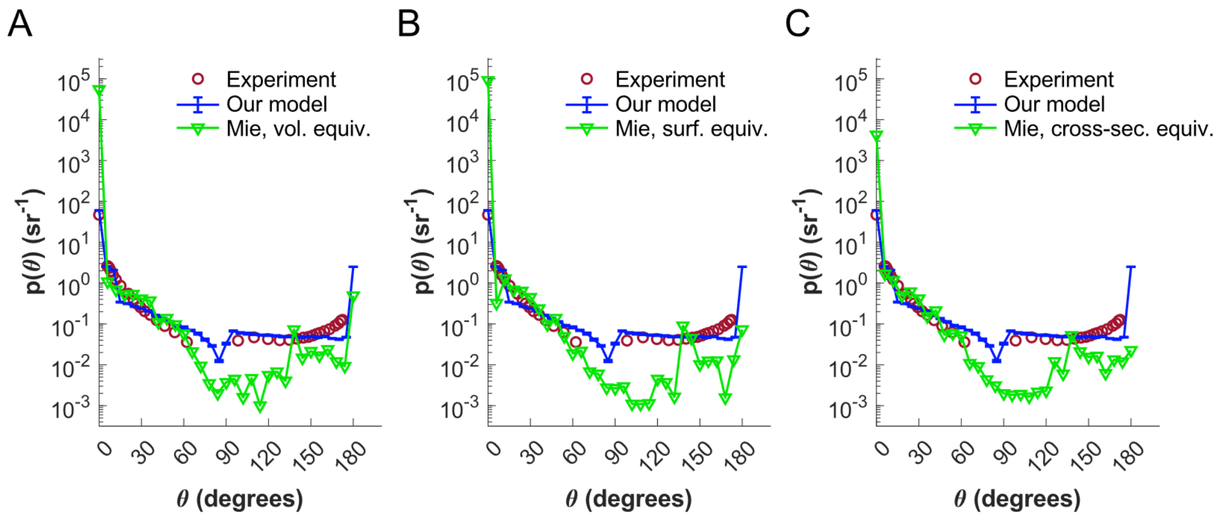


**Fig. S3.** Scattering functions of day-1 adult N2 *C. elegans* (N = 3 to 5 animals) obtained from the experiment (on seeded agar under 629-nm illumination), our non-index matched model (N = 10 trials), and the Mie theory.[1] Referencing to the dimensions of adult *C. elegans* (diameter = 50 µm, length = 1000 µm, n = 1.33), the Mie theory describes the scattering from a sphere with (A) equivalent volume (diameter = 155 µm), (B) surface area (diameter = 226 µm), or (C) cross-section area (diameter = 50 µm). Each datapoint represents mean of measurements. Maximum RSE for the experimental data is 9%. The error bars for the data from our model represent standard error.

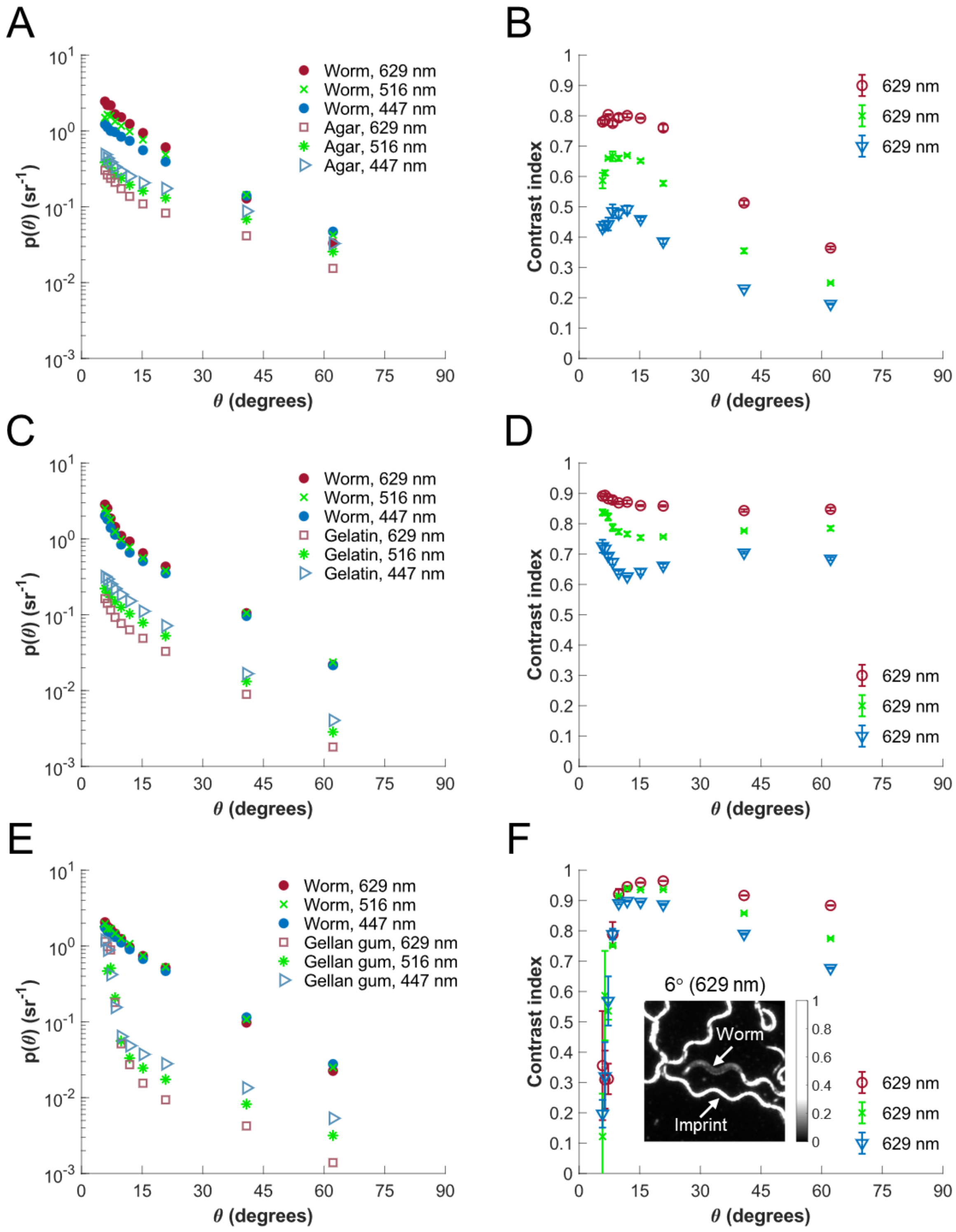


**Fig. S4.** Angle-resolved, wavelength-dependent scattering functions and image contrast for *C. elegans* on unseeded media made with (A and B) agar, (C and D) gelatin, and (E and F) gellan gum. Inset of (F): a representative image of *C. elegans* and its imprints on unseeded gellan gum under a red illumination applied at 6º (FOV: 2.22 × 2.22 mm). For both (A) and (C), maximum RSE is 9% for all the categories. For (E), maximum RSE is 8% for worm and 46% for gellan gum. For (A), (C), and (E), error bars are not shown due to the high density of the datapoints, and statistical details are provided in Table S1. For (B), (D), and (F), error bars represent standard error. For all the panels, each data point represents mean of measurements across N = 3 to 7 animals. All the animals are *C. elegans* N2 at day-1 adulthood.
